# Mind the gap: Nodes of Ranvier are remodeled by chronic psychosocial stress and neuronal activity

**DOI:** 10.1101/2022.04.05.487090

**Authors:** M-K Koskinen, MA Laine, A Abdollahzadeh, A Gigliotta, G Mazzini, SH Journée, V Alenius, K Trontti, J Tohka, P Hyytiä, A Sierra, I Hovatta

**Author notes:** equal contribution (M-K Koskinen, MA Laine). Correspondence should be addressed to: Prof. Iiris Hovatta, SleepWell Research Program, P.O. box 21, 00014 University of Helsinki, Finland.

## Abstract

Differential expression of myelin-related genes and changes in myelin thickness have been demonstrated in mice after chronic psychosocial stress, a risk factor for anxiety disorders. To determine whether and how stress affects structural remodeling of nodes of Ranvier, another form of myelin plasticity, we developed a 3D reconstruction analysis of node morphology in C57BL/6NCrl and DBA/2NCrl mice. We identified strain-dependent effects of chronic stress on node morphology, including elongation of paranodes in the medial prefrontal cortex (mPFC) in DBA/2NCrl mice. Furthermore, chronic chemogenetic activation of the ventral hippocampus-to-mPFC pathway resulted in increased risk assessment behavior and shortened paranodes specifically in stimulated axons, providing a direct link between anxiety-like behavior and remodeling of the nodes. Altogether, our data demonstrate genetic regulation of nodal remodeling in stress and suggest an activity-dependent regulation of paranodes in anxiety-related circuits. Nodal remodeling may thus contribute to the aberrant circuit function associated with anxiety disorders.

## Introduction

Anxiety disorders are the most common psychiatric disorders, affecting up to 14% of the population^1^, with an immense socioeconomic burden^2^. Anxiety disorders are complex diseases with both genetic and environmental risk factors^3,4^, however, little is known about the mechanisms underlying these gene-environment interactions. Chronic psychosocial stress increases the risk to develop anxiety disorders^5,6^. Understanding the mechanisms mediating vulnerability to stress, and resilience against it, is crucial as it could provide much-needed treatment and prevention strategies.

By employing the chronic social defeat stress (CSDS) model, a well-validated animal model of psychosocial stress and anxiety-like behavior in male mice^7,8^, we recently demonstrated using four inbred mouse strains that behavioral responses to stress are strongly influenced by genetic background^9^. CSDS, consisting of confrontations between intruder and resident mice, leads to social avoidance in a subset of mice (stress-susceptible), while others retain social approach (stress-resilient) similar to non-stressed mice. For example, most (~95%) DBA/2NCrl (D2) mice show social avoidance after CSDS, and are thus classified as stress-susceptible, while the majority (~70%) of C57BL/6NCrl (B6) mice are stress-resilient^9^.

Our prior unbiased gene expression profiling study, aimed to identify biological pathways mediating stress-induced anxiety and resilience to it, discovered statistical over-representation of myelin-related genes among differentially expressed genes in mice after CSDS^9^. Both the genetic background, as well as resilience and susceptibility to stress modulated this effect. Changes in gene expression were accompanied by strain- and susceptibility/resilience-dependent differences in myelin sheath thickness in the medial prefrontal cortex (mPFC), ventral hippocampus (vHPC), and bed nucleus of stria terminalis (BNST), regions involved in the regulation of anxiety^10^. Altogether, these findings, also supported by others^11–15^, highlight the involvement of myelin plasticity in stress response and suggest an important role for myelination in mediating susceptibility and resilience to stress.

In addition to changes in myelin thickness, myelin plasticity involves modulation of the nodes of Ranvier^16^, small unmyelinated segments between internodes that enable saltatory conduction^17^. Nodes of Ranvier are characterized by unmyelinated nodes containing voltage-gated sodium ion channels. These node gaps are flanked by paranodes, where myelin sheaths attach to axons, and by adjacent juxtaparanodes that contain a high density of potassium channels^17–19^. Similarly to changes in myelin thickness, modulation of nodes of Ranvier properties can have profound changes on axonal conduction^20–23^, and thus critically impact connectivity of neuronal networks^24^. Consequently, disruption of the nodes of Ranvier, as observed for example in neurodegenerative disorders^25–27^, may contribute to aberrant axonal conduction evident in these conditions.

Here, we investigated whether modulation of the nodes of Ranvier structure occur in response to CSDS, and whether these changes are influenced by genetic background or resilience or susceptibility to stress-induced anxiety. We identify strain-dependent effects of chronic stress on nodal structures using a novel 3D segmentation method and show that these changes occur differently in the mPFC gray and white matter. Importantly, we also demonstrate that projection-specific modulation of neuronal activity by DREADDs (Designer Receptors Exclusively Activated by Designer Drugs)^28^ in the vHPC-to-mPFC pathway, that led to changes in anxiety-like behavior, induced axon-specific differences in paranode length. Altogether, we identify node of Ranvier plasticity as a key phenomenon involved in stress response, and its dependence on genetic background. Furthermore, we demonstrate that neuronal activity drives paranode modifications in an axon-specific manner in anxiety-related brain circuits.

## Results

### Node of Ranvier-related genes are differentially expressed in response to chronic psychosocial stress

In our previous study, we discovered changes in the expression of myelin-related genes in response to CSDS^9^. These changes were accompanied by differences in myelin thickness, which occurred in a brain region- and strain-dependent manner, altogether demonstrating myelin plasticity as an integral part of the stress response^9^. These findings prompted us to further investigate whether chronic stress impacts other myelin features, such as the nodes of Ranvier. To investigate differential expression of genes involved in the structure and function of nodes of Ranvier (**Figure 1a**, Supplementary table 1), we took advantage of our previously published RNA-sequencing dataset from the medial prefrontal cortex (mPFC) of B6 and D2 mice, performed 1 week after cessation of a 10-day CSDS^9^. Amongst the 40 genes examined, we found that 12 genes (30%) were differentially expressed in B6 susceptible animals compared to controls, with 11 of them (28%) being expressed at a lower level in the susceptible compared to control mice (**Figure 1b**). These genes encoded for voltage-dependent ion channels (e.g., *Kcnq2*; Kv7.2 and *Scn1a*; Nav1.1) and structural components of the nodes (e.g., *Gjc*; Connexin-47; *Mag;*Myelin associated glycoprotein). In addition, to examine potential statistical enrichment of the genes associated with the structural subcomponents of the nodes of Ranvier (**Figure 1a**), we conducted Gene Set Enrichment Analysis (GSEA)^29^ on differential gene expression lists of B6 resilient, B6 susceptible, and D2 susceptible mice compared to controls (**Figure 1c&d**). GSEA tests a statistical hypothesis of a gene set being overrepresented among the up- or downregulated genes. We found that paranodal, juxtaparanodal, and nodal genes were significantly overrepresented among the downregulated genes in the D2 susceptible mice compared to same-strain controls. In B6 susceptible animals, a statistical trend for overrepresentation of nodes of Ranvier genes among the downregulated genes was observed. We found similar results when GSEA was performed with all 40 genes associated with nodes of Ranvier (**Figure 1c&d**). Taken together, these results suggest that psychosocial stress alters the expression levels of genes involved in nodes of Ranvier structure and function in the mPFC of B6 and D2 susceptible animals.

**Figure 1.**
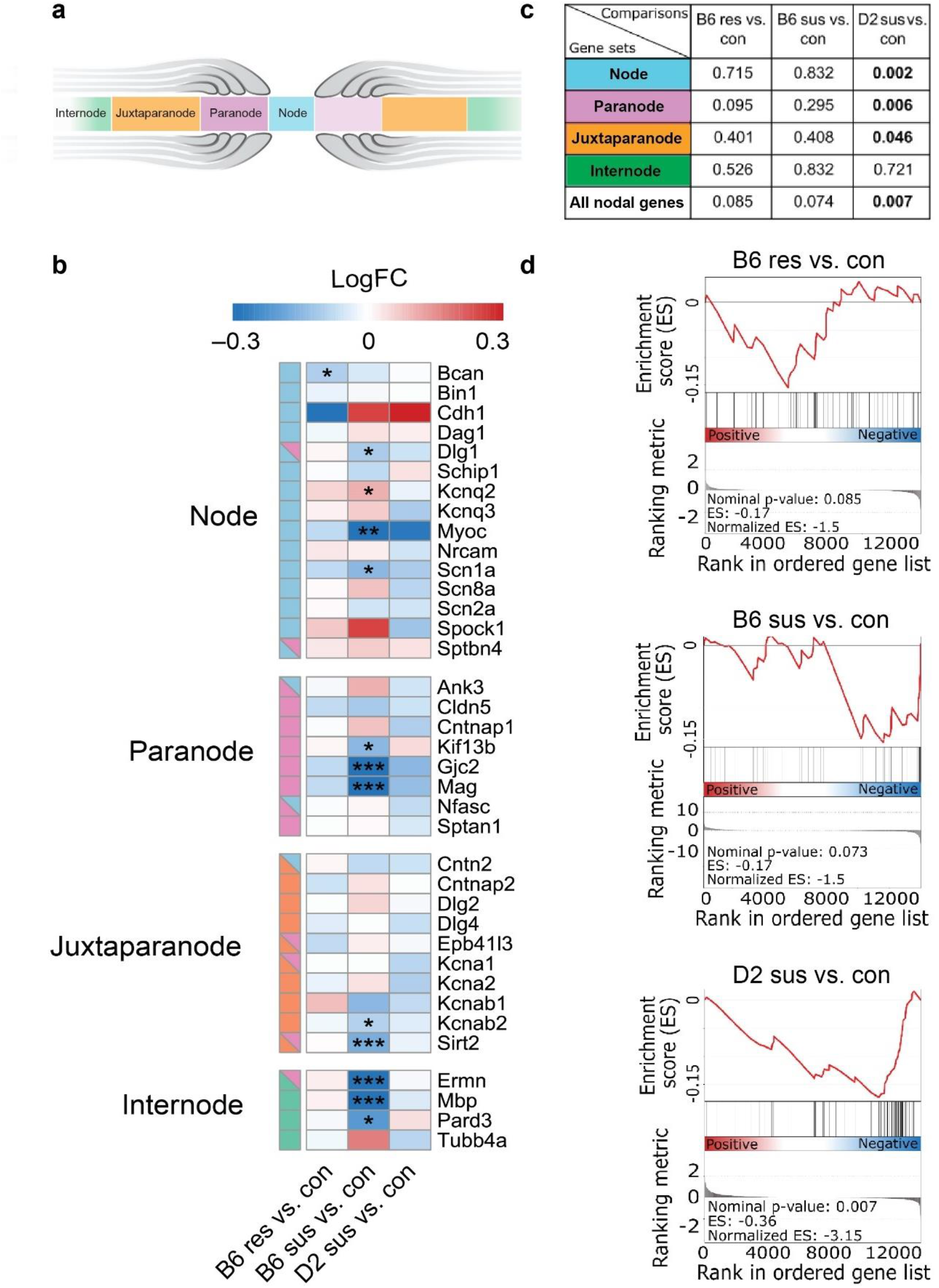
Differential expression of nodes of Ranvier genes after chronic social defeat stress. **a** Schematic representation of the node of Ranvier subregions. **b** Heatmap showing the expression fold change (logFC) and significance of differential expression for genes associated with nodes of Ranvier subcomponents in B6 and D2 defeated mice in the mPFC. **c** Gene Set Enrichment Analysis (GSEA) of genes associated with the individual structural subcomponents of the nodes of Ranvier (node, paranode, juxtaparanode, internode) and combined analysis of all nodal genes (N=40 genes, see supplementary table 1). **d-f** GSEA enrichment score figures of all nodal genes. The top portion of the plot shows the enrichment score, which reflects the degree of overrepresentation of the gene set at the top or bottom of a ranked gene list. The middle portion of the plot shows the position of the genes in the ranked list. The bottom portion of the plot shows the value of the ranking metric as the analysis walks down the list of ranked genes. B6: C57BL/6NCrl; D2: DBA/2NCrl; Con: control; Res: resilient; Sus: susceptible.

### Strain-dependent alterations of nodal morphology after chronic psychosocial stress

To investigate whether the observed gene expression differences associate with changes in nodes of Ranvier morphology, we developed a novel 3D reconstruction analysis of nodal subdomains, including the paranode and juxtaparanode sections. We analyzed these nodal subdomains after CSDS in the anterior cingulate cortex subregion of the mPFC, a critical hub for the regulation of anxiety^30^. This brain region contains several short- and long-range connections^31^ as well as excitatory and inhibitory neurons, both of which are known to be myelinated^32^. To investigate whether genetic background or susceptibility or resilience to stress influence nodal modifications in response to stress, we conducted the morphological analysis both in B6 and D2 mice, classified either as stress-resilient or -susceptible based on social avoidance behavior following CSDS^9,33^ (Supplementary **Figure 1**).

We found that D2 mice had longer paranodes, identified by contactin-associated protein (CASPR) immunoreactivity (**Figure 2a&b**), both in resilient (by 9.4%) and susceptible (by 10.7%) mice compared to same strain controls (**Figure 2c**). Conversely, paranode length did not differ in stressed B6 mice compared to controls, yet we found that paranode length was increased (6%) in susceptible mice compared to same strain resilient mice. Moreover, node width, defined as the distance between the two flanking paranodes (**Figure 2a)**, was shorter (6.6%) in B6 susceptible mice compared to same strain resilient mice, while unaffected in D2 mice (**Figure 2d**). In accordance with paranodal elongation, total node length was longer in D2 resilient mice (by 10%) compared to same strain controls (**Figure 2e**).

**Figure 2.**
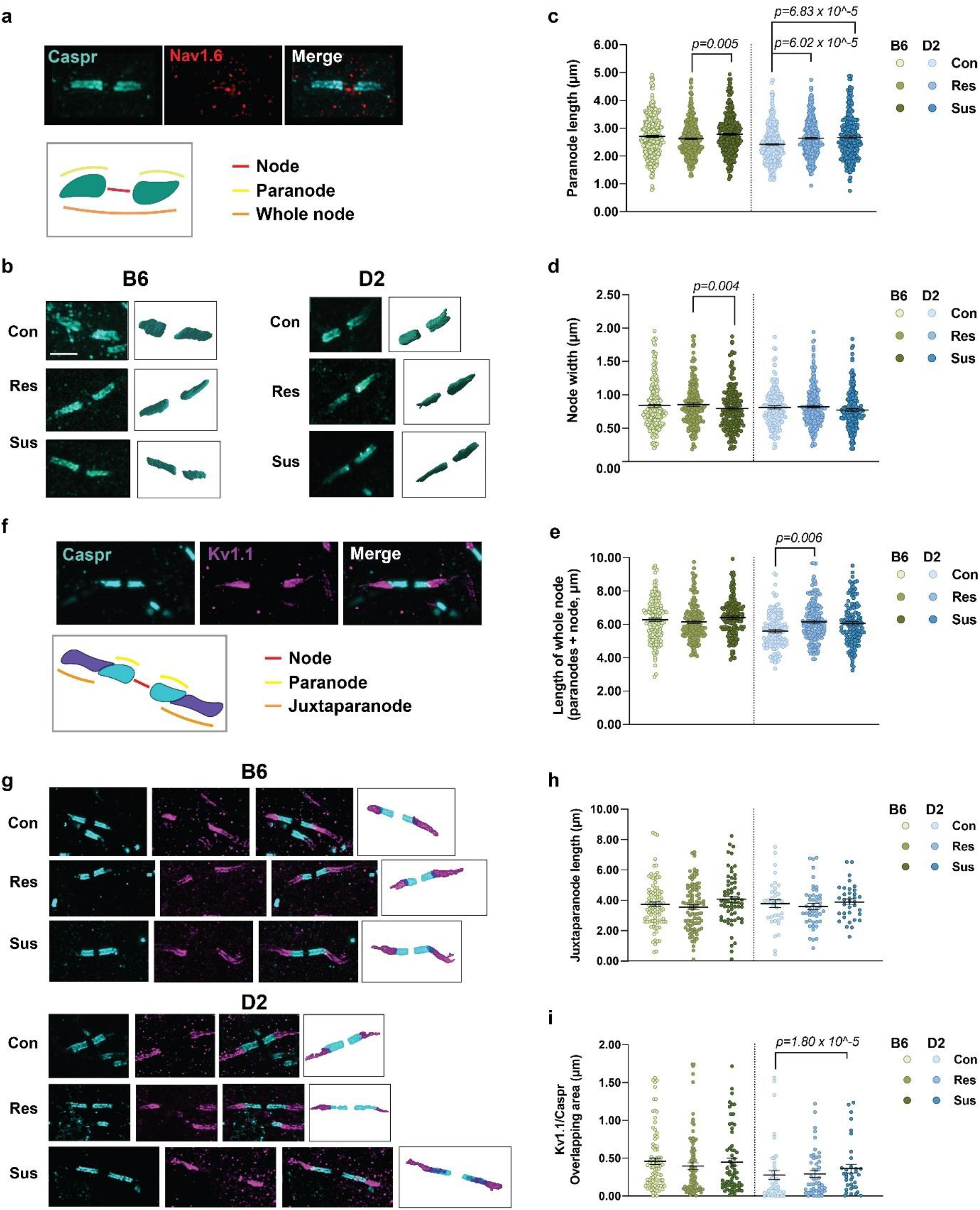
Strain-dependent changes in nodal morphology after chronic social defeat stress. **a,b** 3D reconstructed paranodes in the anterior cingulate cortex layer V/VI in B6 and D2 mice after chronic social defeat stress. **c-e** Quantification of paranode length (**c**), node width (**d**) and total node length (**e**) (**c** Paranode length B6: Con=349 paranodes from 5 mice, Res=345 paranodes from 5 mice, Sus=357 paranodes from 5 mice; Generalized estimating equation (GEE) *p*=0.018, pairwise comparisons: Con *vs*. Res, *p*=0.155; Con *vs*. Sus, *p*=0.208; Res *vs*. Sus, *p*=0.005. D2: Con=308 paranodes from 5 mice, Res=372 paranodes from 5 mice, Sus=296 paranodes from 5 mice, GEE *p*=1.86×10^-5^, Con *vs*. Res, *p*=6.02×10^-5^; Con *vs*. Sus, *p*=6.86×10^-5^; Res *vs*. Sus, *p*=0.601. **d** Node width B6: Con=201 nodes from 5 mice, Res=193 nodes from 5 mice, Sus=187 nodes from 5 mice, GEE *p*=0.013, Con *vs*. Res, *p*=0.718; Con *vs*. Sus *p*=0.237; Res *vs*. Sus *p*=0.004. D2: Con=164 nodes from 5 mice, Res=220 nodes from 5 mice, Sus=200 nodes from 6 mice; GEE *p*=0.316. **e** Total node: B6: Con=157 nodes from 5 mice, Res=157 nodes from 5 mice, Sus=167 nodes from 5 mice; GEE *p*=0.367. D2: Con=131 nodes from 5 mice, Res=172 nodes from 5 mice, Sus=129 nodes from 5 mice; GEE *p*=0.022, Con *vs*. Res, *p*=0.006; Con *vs*. Sus *p*=0.097; Res *vs*. Sus *p*=0.683). **f,g** 3D reconstructed juxtaparanodes and the overlapping region between paranodal and juxtaparanodal sections. **h,i** Quantification of juxtaparanode length and (**i**) Caspr /Kv1.1 overlapping area (**h**Juxtaparanode length: B6: Con=92 nodes from 4 mice, Res=92 nodes from 4 mice, Sus=64 nodes from 3 mice; GEE *p*=0.472. D2: Con=36 nodes from 4 mice, Res=47 nodes from 3 mice, Sus=35 nodes from 3 mice; GEE *p*=0.211. **i** Overlapping area: B6: Con=89 nodes from 4 mice, Res=91 nodes from 4 mice, Sus=65 nodes from 3 mice; GEE *p*=0.472. D2: Con=43 nodes from 4 mice, Res=48 nodes from 3 mice, Sus=35 nodes from 3 mice; GEE *p*=0.001, Con *vs*. Res, *p*=0.457; Con *vs*. Sus *p*=1.80×10^-5^; Res *vs*. Sus *p*=0.007). B6: C57BL/6NCrl; D2: DBA/2NCrl; Con: control; Res: resilient; Sus: Susceptible. All nominal *p*-values surviving Bonferroni correction are shown. Error bars represent ± SEM.

Changes in paranode length have been associated with disruption of paranodal junctions that may result in disorganization of nodal subdomains and aberrant localization of juxtaparanodal Kv1-channels to paranodal sites^34–36^. To investigate whether CSDS-associated elongation of paranodes, as observed in D2 mice, co-occur with juxtaparanodal alterations, we assessed morphology of juxtaparanodes, and the overlap of paranodes and juxtaparanodes by employing our 3D segmentation analysis (**Figure 2f**). Juxtaparanode length, identified by Kv1.1 potassium channel immunoreactivity (**Figure 2g**), was not affected by CSDS in either B6 or D2 strain (**Figure 2h**). However, the overlapping area between paranodes and juxtaparanodes, identified as double immunoreactivity of CASPR and Kv1.1, was longer (29%) in D2 susceptible mice compared to same strain controls (**Figure 2i**). Altogether, these findings suggest that although stress does not alter overall juxtaparanodal morphology, it induces mislocalization of juxtaparanodal Kv1.1 channels into paranodal sections in susceptible D2 mice, as observed in aging ^26^ and in multiple sclerosis^25^.

In addition to cortical gray matter, we analyzed nodal morphology in the forceps minor (Supplementary **Figure 2a**), a white matter tract adjacent to the mPFC. Paranode length did not change in response to CSDS in the white matter in either B6 or in D2 mice (Supplementary **Figure 2b**). Analysis of nodal width, however, demonstrated shorter (10.7%) node gap in stress susceptible D2 mice compared to same strain resilient mice, and no differences between the groups in the B6 strain (Supplementary **Figure 2c**). Altogether, our analysis demonstrates that chronic stress alters node of Ranvier morphology both in the mPFC gray and white matter. Importantly, these effects are strongly influenced by genetic background and depend on the individual stress response.

### No differences in the number of oligodendrocyte lineage cells after chronic psychosocial stress

Differences in nodes of Ranvier length could be influenced by addition of new oligodendrocytes (OLG), or lack thereof^37^. To address whether CSDS altered the number of OLGs or oligodendrocyte progenitor cells (OPC), the source of differentiated OLGs, we quantified the density of these cell populations in the mPFC. We did not detect differences in the number of OPCs or OLGs after CSDS in either mouse strain (Supplementary **Figure 3**), suggesting that nodal changes involve modification of existing myelin sheaths.

### Repeated activation of the vHPC-mPFC pathway decreases anxiety-like behavior and reduces paranode length in axon-specific manner

As neuronal activity is a major regulator of myelin plasticity^38–40^, and transcranial magnetic stimulation affects node width^21^, we hypothesized that nodal changes, as observed after CSDS, arise due to stress-induced activity changes in brain circuits affected by stress^41^. To determine whether nodal alterations are directly driven by changes in neuronal activity, we chemogenetically activated neurons and studied nodal changes along the axons of these cells. For this, we chose a pathway projecting from the vHPC to the mPFC, previously shown to regulate anxiety-like behavior^42,43^, and expressed the excitatory hM3Dq DREADD in these projection neurons.

To achieve projection-specific expression of hM3Dq-receptors, we bilaterally injected a retrograde *Cre*-carrying virus (AAV_retro_-Cre-eGFP) into the mPFC, while a *Cre-*dependent hM3Dq (AAV8-Dio-hM3Dq-mCherry) or an mCherry (AAV8-Dio-mCherry) control vector was injected into the vHPC (Figure **3a**) of male C57BL/6NCrl mice. Immunohistochemical analysis confirmed the accuracy of the labeling by showing robust eGFP labeling in the mPFC and mCherry expression in the vHPC CA3/1 regions, as well as eGFP/mCherry-double positive axons along the pathway (Figure **3b**).

**Figure 3.**
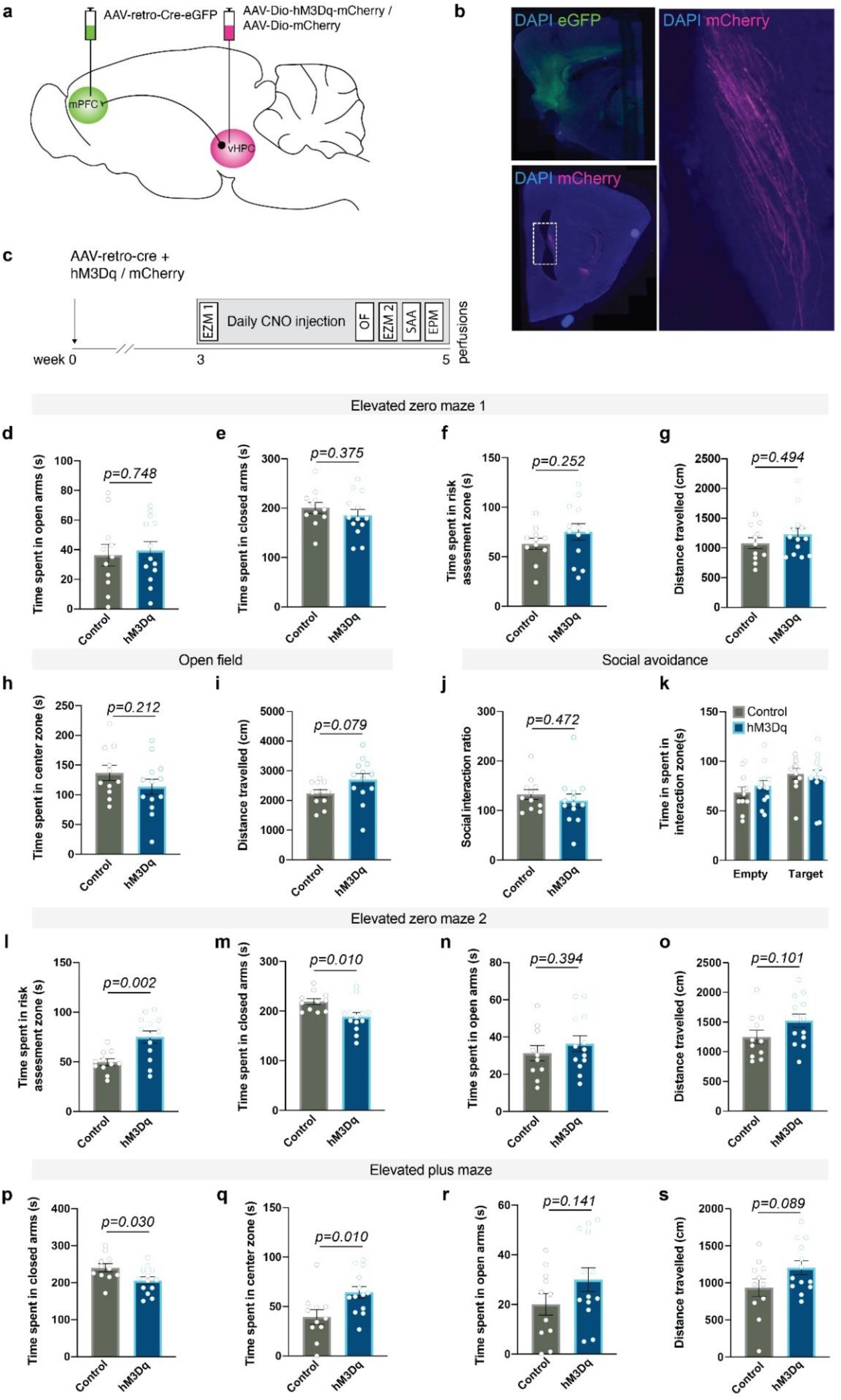
DREADD-mediated activation of vHPC-to-mPFC projection neurons reduces anxiety-like behavior. **a** Projection-specific expression of hM3Dq/mCherry was achieved by injecting a retrograde *Cre*-carrying virus (retro-Cre-eGFP) into the mPFC and a *Cre*-dependent hM3Dq (Dio-hM3Dq-mCherry) or a control virus (Dio-mCherry) into the vHPC CA3 subregion. **b** Representative examples showing eGFP expression in the mPFC and mCherry-expressing axons in the hippocampal fimbria. **c** Experimental timeline. **d-f** Closed area, (**e**) open area and (**f**) risk assessment (RA) zone time in the acute elevated zero maze test (EZM1) (two-sided student’s t-tests). **g** Distance travelled during the EZM1 test (two-sided unpaired Mann-Whitney test). **h-i** Time spent in the center of the open field (OF) test and (**i**) Distance travelled during the OF test (two-sided student’s t-tests). **j** Quantification of social interaction ratio in the social approach-avoidance (SAA) task (two-sided student’s t-test). **k** Time spent in the interaction zone during empty and target sessions of the SAA test (Two-way repeated ANOVA: group x session F(1,22)=0.703, *p*=0.402; session F(1,22)=6.111, *p*=0.022; group F(1,22)=0.100, *p*=0.755). **l-n** Time in the RA zone (two-sided Welch’s t-test), (**m**) closed area and (**n**) open area (two-sided student’s t-tests) in the EZM2 test after repeated activation of vHPC-to-mPFC pathway. **o** Distance travelled during the EZM2 test (two-sided student’s t-test). **p-r** Time spent in the closed arms, (**q**) center zone and in (**r**) the open arms in the elevated plus maze (EPM) test (two-sided student’s t-tests). **s** Distance travelled in the EPM test (two-sided student’s t-test). hM3Dq n=13; Control n=11. Error bars represent ±SEM.

Behavioral tests confirmed that the activation of this specific pathway was effective in modulating anxiety-like behavior (Figure **3c**). We first assessed the effects of an acute chemogenetic activation of Gq signaling on anxiety-like behavior by testing the mice in the elevated zero maze (EZM) 20-30 minutes after a single clozapine-*N*-oxide (CNO) injection, which activates the hM3Dq receptors. In the EZM, both hM3Dq and control groups spent equal amount of time in the open areas, closed areas, and in the risk assessment zone, and no effect on general activity was observed (**Figure 3d-g**), suggesting ineffectiveness of the acute manipulation to alter anxiety-like behavior in this assay. After the acute test, we continued daily CNO injections over several days to assess the effects of a repeated manipulation of vHPC-to-mPFC activity on anxiety-like behavior. After 12 days of CNO injections, we carried out the open field (OF) test. No differences in anxiety-like behavior or in general activity were observed between the groups, as demonstrated by similar time spent in the center zone and the distance travelled (**Figure 3h-i**). On the following day, and after another CNO injection, we re-tested the mice in the EZM task. In contrast to the acute CNO effects, after the repeated manipulation hM3Dq mice spent more time in the risk-assessment zone, compared to controls (**Figure 3l**). This was accompanied with reduced closed area time, while the time spent in the open areas or general activity did not differ between groups (**Figure 3m-o**). Next, we assessed the effects of the repeated manipulation on social behavior by testing the mice in the social approach avoidance (SAA) task the following day (after 14 days of CNO injections). hM3Dq and control mice spent equal amounts of time interacting with a social target (**Figure 3j-k**), suggesting that repeated manipulation of the vHPC-to-mPFC pathway activity does not alter social approach-avoidance behavior.

Lastly, we tested the priming effects of an acute hM3Dq activation on anxiety-like behavior. After receiving daily CNO injections for 15 days, the mice received a single CNO injection 30 minutes prior to testing in the elevated plus maze (EPM), to avoid a habituation effect from repeated testing in the EZM. An acute activation in primed animals resulted in reduced exploration of the closed arms in hM3Dq mice, which was accompanied with increased center zone time compared to controls, with no change in open arm time or in locomotor activity (**Figure 3p-s**). Taken together, our behavioral data demonstrate that prolonged manipulation of vHPC-to-mPFC activity decreases anxiety-like behavior, reflected particularly as increased risk assessment in the EZM test and reduced closed arm exploration in the EPM test.

Finally, we assessed whether DREADD-mediated activation, changing anxiety-like behavior, altered nodal morphology along labeled vHPC-to-mPFC axons. For this, we imaged paranodes (identified with CASPR) on mCherry-expressing (mCherry+) axons within the hippocampal fimbria, where robust labeling of mPFC-projecting vHPC axons was observed in both hM3Dq and control mice (Figure **4a&b**). To tease out manipulation-specific effects, we also studied paranodes on non-labeled axons (mCherry-) within the same brain region. In mice with vHPC-to-mPFC projection neurons expressing hM3Dq, paranodes were significantly shorter (9.6%) than in controls (**Figure 4c**). Importantly, the effect of our manipulation was specific for paranodes on mCherry-labeled axons, as no difference in paranode length was present between the groups in non-labeled axons. Furthermore, paranode length did not differ between labeled and non-labeled axons in control mice. We did not observe any effect of the manipulation on node width nor on whole node length (**Figure 4d-e**). These results demonstrate that manipulation of neuronal activity specifically affected paranode length, and, importantly, the modification occurred in the activated, but not non-activated, axons, supporting our hypothesis on activity-dependent remodeling of nodes.

**Figure 4.**
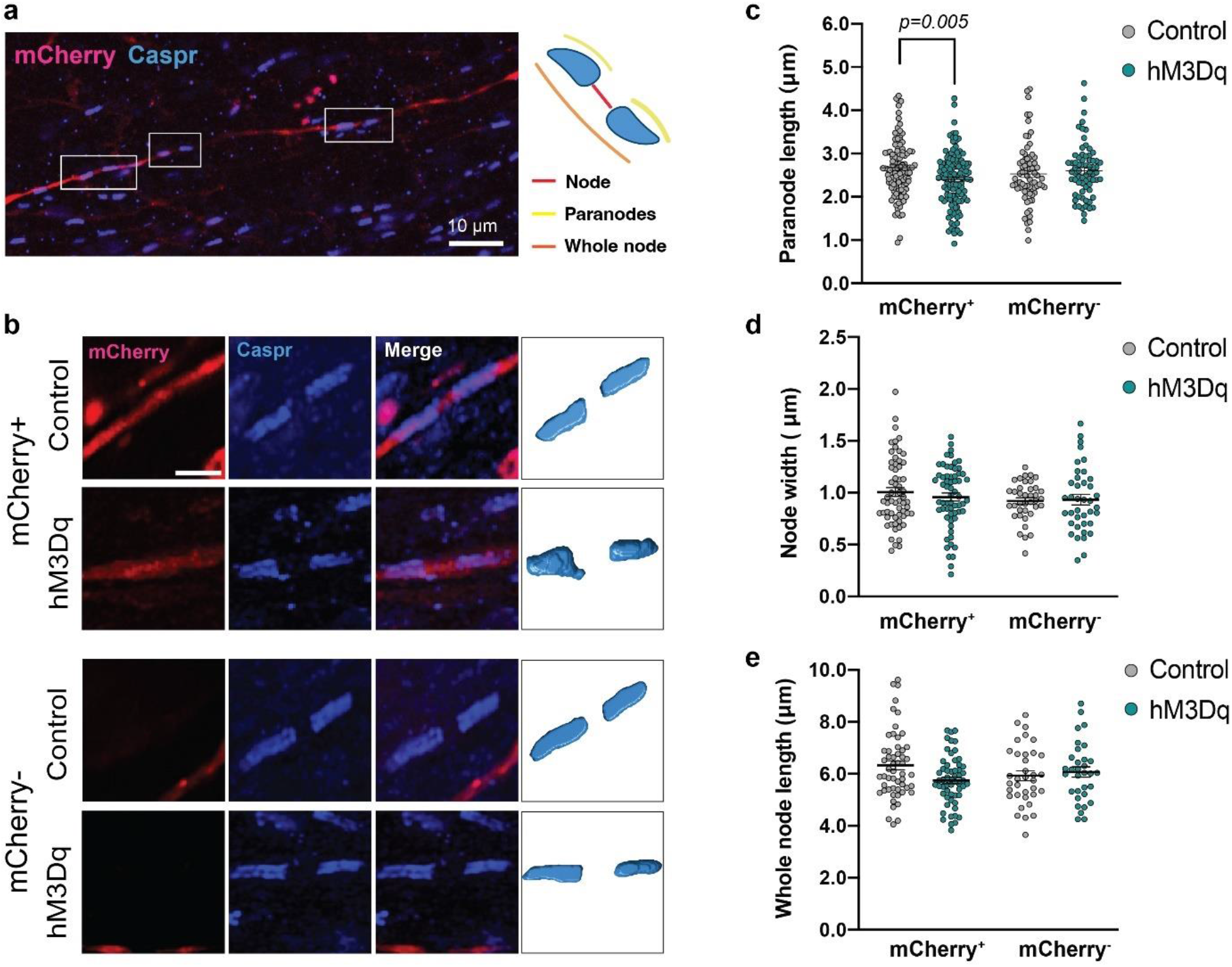
Repeated activation of vHPC-to-mPFC projection neurons reduces paranodal length. **a** A sagittal section containing the vHPC immunolabeled with Caspr antibody in mice expressing an mCherry-labeled hM3Dq or a control virus in vHPC-to-mPFC projection neurons. Outlines demonstrating paranodes along an mCherry-labeled axon. **b** 3D reconstruction of paranodes. **c-e** Quantification of paranode length (Generalized estimating equation: Group x Axon-type, *p*=0.036; pairwise comparisons: hM3Dq-mCherry^+^ vs. Control-mCherry^+^, *p*=0.005; hM3Dq-mCherry^-^ vs. Control-mCherry^-^, *p*=0.420; hM3Dq-mCherry^+^ vs. hM3Dq-mCherry^-^, *p*=0.075; Control-mCherry^+^ vs. Control-mCherry^-^, *p*=0.273), (**d**) node width (Group x Axon-type, *p*=0.205) and (**e**) total node length (Group x Axon-type, *p*=0.261) after repeated activation of the vHPC-to-mPFC pathway. (**c** hM3Dq: 70 mCherry- and 122 mCherry+ paranodes from 6 animals; Control: 72 mCherry- and 122 mCherry+ paranodes from 6 animals. **d** hM3Dq: 39 mCherry- and 62 mCherry+ paranodes from 6 animals; Control: 38 mCherry- and 64 mCherry+ paranodes from 6 animals. **e** hM3Dq: 38 mCherry- and 57 mCherry+ paranodes from 6 animals; Control: 34 mCherry- and 54 mCherry+ paranodes from 6 animals). **b** Size bar 2 μm. Error bars represent ±SEM. Nominal p-values surviving Bonferroni correction are shown.

## Discussion

In this report, we investigated the morphology of nodes of Ranvier after chronic psychosocial stress. These studies were based on our initial finding of differential expression of nodes of Ranvier-related genes within the mPFC of mice exposed to CSDS. The observed gene expression differences were accompanied by altered node of Ranvier morphology. Furthermore, we assessed whether altered neuronal activity produces nodal changes and showed shortened paranodes along DREADD-stimulated axons in a pathway that regulates anxiety-like behavior. Our data provides a proof-of-concept that chronic activity can alter node of Ranvier morphology along the active axons, and that such changes can be accompanied by measurable changes in behavior.

We found longer paranodes in chronically stressed D2 mice compared to controls in the mPFC gray matter. Similarly, in B6 mice paranodes were longer in stress-susceptible mice compared to resilient mice. Elongated paranodes may indicate loosening of paranodal junctions, as observed for example in neurodegenerative disorders^25,44^ and in aging^26,45^. Loosening of myelin-axon contacts at the paranodes may result in separation of the paranodal loops, which in turn may lead to exposure of juxtaparanodal Kv1 channels^25^ and leakage of these channels to the paranodal sites, all of which can alter axonal conduction and neuronal excitability^34,46,47^. In addition to elongated paranodes, we observed that the border between paranodal and juxtaparanodal domains was disrupted in stress-susceptible D2 mice compared to same strain control mice. This could indicate a leakage of juxtaparanodal Kv1.1 channels to paranodal sites, and thus disruption of axonal conduction.

Myelination in both the white and gray matter contribute to experience-dependent shaping of neural circuits^16,40^. Therefore, we also assessed nodal remodeling in the white matter after chronic stress and found shortened nodes in stress-susceptible D2 mice compared to same-strain resilient mice within forceps minor, the anterior part of the corpus callosum. A previous study assessing the effects of chronic restraint stress on the nodes of Ranvier morphology in B6 mice demonstrated shorter nodes and longer paranodes in the corpus callosum^48^. We did not, however, observe paranodal changes within forceps minor in either strain. This discrepancy could result from methodological differences, such as type and duration of the stressors and analysis method. Taken together, our data demonstrate different effects of CSDS on the nodes of Ranvier in the mPFC gray and white matter, suggesting regionally specific impact of stress. As morphological differences were accompanied by differential expression of node of Ranvier-related genes, it is plausible that remodeling occurs via active, cell-dependent mechanisms. Furthermore, the number of OPCs and OLGs remained unaltered, at least as assessed one week after cessation of stress, suggesting that nodal plasticity may involve modification of existing myelin, as observed after repeated neuronal stimulation and learning^21^.

What drives the remodeling of the nodes of Ranvier after chronic stress? We set out to answer this question by using a chemogenetic approach in the vHPC-to-mPFC pathway, previously shown to regulate anxiety-like behavior^42,43^. We found that prolonged DREADD-mediated activation of vHPC-to-mPFC projection neurons resulted in increased risk-assessment and reduced closed arm exploration, suggesting decreased anxiety-like behavior. An acute inhibition of the pathway has previously been shown to decrease anxiety-like behavior^42,43^, while an acute activation elicited an anxiogenic effect^43^. Altogether, these data demonstrate that vHPC-to-mPFC pathway is critical for the regulation of anxiety-like behavior, yet it underscores a divergent impact of an acute and chronic activation of the pathway on anxiety. We next asked whether nodal changes occur in an activity-dependent manner in the vHPC-to-mPFC pathway.

Whether activity-related nodal alterations result from generalized activity changes within a brain region, or whether nodal modulation occurs at the level of individual axons, has remained unanswered. We used chemogenetics because this method allowed us to repeatedly stimulate specific neurons, mimicking a chronic stress situation, while visualizing the manipulated axons for morphological analysis of nodes. Notably, paranodes were specifically modified on activated axons, but not on non-activated axons within the same brain region, suggesting that neuronal activity directly elicits paranodal plasticity along the stimulated axons, as described previously for the formation of myelin sheaths^39^. Axon-specific modulation of nodes of Ranvier likely adds to fine-tuning of axonal conduction, enabling circuit-specific adaptation to changing processing needs. However, it could also underlie some of the persistent maladaptations associated with stress-related psychiatric disorders.

We found that the main nodal effect of CSDS was lengthening of paranodes, while DREADD-mediated stimulation of axons resulted in shortening of paranodes. We have previously shown that chronic stress can associate both with thicker or thinner myelin sheaths, depending on whether the mice are stress-susceptible or -resilient and on the genetic background^9^. Similar to dynamic modulation of myelin sheaths^40^, bidirectional modulation of paranodes likely contributes to the capacity to adapt according to the needs of specific circuits and networks^20^.

Overall, our data demonstrate genetically controlled remodeling of nodes of Ranvier after chronic stress. Much of the past work on myelin plasticity and stress has been conducted using only one mouse strain, typically a B6 substrain. Our findings highlight the need to study individuals from more than one genetic background, as appreciating genetically dependent outcomes could improve translational validity of preclinical models. We propose that nodes of Ranvier remodeling associated with CSDS result from stress-induced changes in neuronal activity patterns in distinct brain regions, including the mPFC^41^. Similar to modification of myelin sheaths^40^, nodal changes altering processing in affected circuits may either promote susceptibility or resilience to stress. We cannot, based on our data, characterize the observed modifications as pathological or adaptive. However, the similarity of our results to those related to senescence^26^ and multiple sclerosis^25^ suggest pathology of the CSDS-associated changes in stress-susceptible mice. As anxiety disorders are increasingly considered to involve circuit-level dysfunction, rather than a disruption of a single neurotransmitter system or a single brain region^49^, nodal remodeling may contribute to aberrant circuit function in these conditions. Future studies should be aimed not only to understand how nodes of Ranvier remodeling influences conduction velocity along single axons, but to understand its role in circuit- and network level communication. Understanding the mechanisms driving node of Ranvier changes may help pave the way for novel treatment practices for anxiety disorders.

## Materials and methods

### Animals

C57BL6/NCrl (B6) and DBA/2NCrl (D2) male mice (5 weeks old upon arrival, Charles River Laboratories) were purchased for all experiments and allowed to acclimatize for 10 (Experiment 1: CSDS) or 7 (Experiment 2: DREADD) days before the start of procedures. As aggressor mice in CSDS we acquired Crl:CD1 (CD1, Charles River Laboratories) male mice aged 13-26 weeks. Mice were initially housed in groups (B6 and D2) or individually (CD1) before CSDS at 22 ± 2 °C and humidity 50 ± 15 % on a 12h light/dark cycle (lights on 6:00 - 18:00). After CSDS and stereotactic surgeries all mice were single-housed. They had ad libitum access to water and food except during behavioral experiments. All animal procedures were approved by the Regional State Administration Agency for Southern Finland (ESAVI/2766/04.10.07/2014 and ESAVI/9056/2020) and conducted in accordance with directive 2010/63/EU of the European Parliament and of the Council.

### Experiment 1: CSDS

All CSDS behavioral procedures were carried out at the end of the light phase. CD1 mice were first screened for appropriate aggressive behavior during three consecutive days. They had to attack a screener mouse (B6 or D2) on at least two consecutive sessions, within a latency interval of 5-90 s. CSDS was then performed as previously described^33,50^. For CSDS, B6 or D2 mice were introduced into the resident aggressor’s compartment for max. 10 min. They were then moved to the other side of the cage, separated from the CD1 by a perforated Plexiglas wall, for the rest of the 24 h. Exposure to CSDS was repeated for 10 days and test mice were subjected to a new aggressor each day. Direct physical contact time was reduced in case of injury. Control mice were housed in pairs in similar cages with no physical confrontation, and cage-mates were changed daily. One day after the end of CSDS, having separated the mice into individual cages, test and control mice were tested for social avoidance (SA). We brought all animals into the testing room 30 min before the test. For the first (no-target) trial, we placed the mouse in the middle of an open arena with an empty Plexiglas cylinder on one side of the cage. The mouse’s movements were tracked with Ethovision XT10 (Noldus Information Technology) for 150 s. The mouse was then returned to the home cage, and the arena cleaned. Immediate after, for the social target trial, the same mouse was placed back into the arena for 150 s, with an unfamiliar CD1 in the cylinder. Time in the interaction zone (IZ) was measured for each trial. A social interaction ratio (time spent in the IZ with the social target present / time spent in the IZ with no target present, multiplied by 100) was calculated for each mouse. For CSDS-exposed animals, susceptible mice were defined as having SI ratios below a boundary defined as the strain-specific control mean score minus one standard deviation^50^. Other CSDS-exposed mice were considered resilient because their social interaction ratio was similar to controls.

### RNA-Sequencing and differential gene expression analysis

We re-analyzed RNA sequencing data from brain samples of B6 and D2 mice after CDSD, published by us previously^50^ (GEO accession GSE109315). Briefly, mice were sacrificed 6-8 days after the last CDSD session and RNA was extracted with TriReagent (Ambion). Sequencing libraries were prepared with ScriptSeq v2 RNA-seq library preparation kit (Epicentre) and sequencing was performed on NextSeq500 (single-end 96 bp; Illumina). Differential expression analysis on voom normalized^51^ gene expression values were performed using limma eBayes^52,53^, comparing resilient and susceptible mice to their same-strain controls. Here, we conducted gene set expression analysis (GSEA Desktop v4.1.0^29,54^) using the differential expression results published in^50^. For the GSEA, we ranked the differential expression gene lists by interaction of p-value and fold change of the genes (logFC*p-value). We then analyzed these lists for enrichment of genes belonging to Gene Ontology (GO) terms of node of Ranvier (GO:all, N=40), node (GO:0033268, N=15), paranode (GO:0033270, N=11), juxtaparanodes (GO:0044224, N=10) and internode (GO:0033269, N=4).

### Experiment 2: DREADD - Stereotactic viral injections

Mice were anesthetized using 5% isoflurane. Once anesthetized (toe-pinch and tail-pinch reflexes were absent) the mouse was transferred onto a stereotactic frame (Kopf) and maintained on 2% isoflurane anesthesia. The injection coordinates for the mPFC were AP: + 2.22 mm, ML: ± 0.35 mm, DV: −2.1 mm, and for the vHPC AP: - 3.4 mm, ML: ±2.9 mm, DV: −4.5 mm. AAV_retro_-hSyn1-chl-EGFP_2A_iCre-WRE-SV40p(A) (AAV_retro_-Cre) was injected into the mPFC for retrograde transport to vHPC neurons projecting to the mPFC to the injection region. For the vHPC viral construct, mice were randomly assigned to receive either the control virus (AAV8-hSyn1-dlox-mCherry(rev)-dlox-WPRE-hGHp(A) or the DREADD (AAV8-hSyn1-dlox-hM3D(Gq)_mCherry(rev)-dlox-WPRE-hGHp(A). All viral constructs were ordered from the Viral Vector Facility at the University of Zürich and ETH Zürich. For each injection, 0.5 μL of the virus was injected using an automated pump (World Precision Instruments) and a 10 μL microsyringe (Hamilton Co.) over the course of 3 minutes. The needle was left in place for another 3 minutes, and then slowly withdrawn (min. 3 minutes). The order of control and DREADD surgeries were balanced across and within days. For post-surgical analgesia, mice were given 5 mg/kg of carprofen subcutaneously before removing from anesthesia. Following surgery, mice were single housed and permitted to recover for 3-4 weeks before beginning behavioral tests, ensuring viral expression.

### Experiment 2: DREADD – Behavioral testing and CNO injections

All behavioral tests (except for the SA test) were carried out at the start of the light phase. The mice were brought into the test room 30 minutes before each test. Light conditions for each test are detailed below, and a timeline for behavioral experiments can be seen in Figure 3. Cages were changed once a week, but never within 24h before a behavioral test. For each test the order of mice (control and DREADD) were randomized using random number generation, and the experimenters performing experiments were blind to the condition of the animal. Mouse movement was recorded and tracked using Ethovision WT (v13, Noldus Technologies).

1. Effects of acute DREADD activation on the elevated zero maze (EZM1) After acclimatization to the dimly (15 lux) lit room, each mouse received a dose (1 mg/kg) of clozapine-N-oxide (CNO; Abcam, cat. no. ab141704, dissolved in saline) by i.p. injection in their home cage. The elevated zero maze (EZM) test was started 20-30 minutes after this. The mouse was placed into the center of one of two closed sections of a circular maze elevated 40 cm above ground. The total time spent in open and closed areas was recorded over 5 minutes and analyzed using Ethovision XT10 software. Additionally we defined a 5 cm long zone at the intersection of the open and closed zones as risk assessment zones^55^, extending equally (2.5 cm) into the open and closed zones. CNO injections were continued once per day, between 8am – 10am, for a total of 15 days. The injection order was varied by using one of four randomly generated order lists each day, and the mice were weighed every second day to ensure correct dosing and monitor wellbeing.
2. Effects of chronic DREADD activation on anxiety-like behavior (OFT, EZM2) On day 13, prior to receiving CNO, we carried out the open field test (OFT). The light conditions were bright to ensure the anxiogenic nature of the test (290 lux). Each mouse was allowed to explore an arena (50 x 50 cm) for 5 minutes. We defined the center of the arena as 5 cm away from the walls at each point. The time the mice spent in the center vs periphery was computed. After the test, each mouse received an injection of CNO and was returned to their home cage. On day 14, the EZM was repeated with slight modifications (EZM2). To enhance novelty and reduce habituation-induced lack of motivation to explore, we added fresh bedding material to the open zones (changed between each mouse). The apparatus, environmental conditions, and recorded parameters were the same as in EZM1. After the test, the mice received a CNO injection.
3. Effects of chronic DREADD activation on social behavior (SA) To test for effects of chronic vHPC-mPFC activation on social avoidance behavior, we performed the SA test at the end of the light phase of day 15. To avoid all acute effects of CNO, the mice did not receive an injection this morning. After acclimating to the test room, each mouse went through two trials of the SA test similarly as after CSDS (see above). Here, a naïve male wild-type B6 mouse was used as a social target. The time spent in the IZ and the social interaction (SI) ratio were computed.
4. Effects of an acute re-activation of the chronically activated projection on anxiety-like behavior (EPM). To explore whether the chronic activation had affected the acute response to CNO, we performed an additional test of anxiety-like behavior following a priming injection. The morning after the SA test (day 16) each mouse received an injection of CNO in the behavioral test room, and after 20-30 minutes they were tested in an elevated plus maze (EPM). The EPM measures anxiety-like behavior with similar parameters as the EZM, but the novel apparatus was expected to minimize habituation-related lack of exploratory drive. To start the test, mice were placed in the center of the apparatus and allowed to freely explore the two opposing closed arms, and the two opposing open arms. Time spent in each arm type was tracked, along with time spent in the center area (as a proxy for risk assessment behavior).^55^

### Experiment 1-Nodes of Ranvier immunohistochemistry

We anesthetized the mice 6-8 days after CSDS with a lethal dose of pentobarbital (Mebunat Vet 60 mg/ml, Orion Pharma). Mice were then transcardially perfused with 37°C 4% paraformaldehyde (PFA) in PBS. After post-fixation in 4% PFA (24 h, +4°C), we cut the brains into 40 μm coronal sections with a Leica VT-1200S vibratome (Leica Biosystems), stored in cryoprotectant as free-floating sections (−20°C) until staining.

For paranode staining in the mPFC, sections were first incubated in 0.5% H_2_O_2_ in TBS for 10 min in RT. Sections were then mounted and stained overnight in +4°C with a cocktail of rabbit anti-Nav1.6 (1:500, #ASC-009, Alomone labs) and mouse anti-CASPR (1:250, #75-001, Neuromab) in 5% NGS 0.5% TBS-T, for nodal and paranodal region staining, respectively. Secondary antibodies were goat anti-rabbit IgG Alexa Fluor 568 (1:400, #A-11011, ThermoFisher Scientific) and goat anti-mouse IgG Alexa Fluor 488 (1:400, #A28175, ThermoFisher Scientific) in 1% NGS in 0.5 % TBS-T. After the last wash, slides were coverslipped with Vectashield + DAPI mounting medium (#H-1200, Vector Laboratories). Co-staining with paranode and juxtaparanode markers was done as follows. Free floating mPFC sections were rinsed 3 times for 10 min in PBS, followed by blocking (5% NGS, 2.5% BSA, 0.25% Triton X-100) for 1 h in RT and incubation with primary antibodies mouse anti-CASPR (1:500, #75-001, Neuromab) and rabbit anti-Kv1.1 (1:300, #APC-009, Alomone labs) in blocking solution overnight in +4°C. Sections were then rinsed 4 times for 10 min in PBS, followed by secondary antibody incubation with goat anti-mouse Alexa Fluor 555 (1:400, #A-21422, ThermoFisher Scientific) and goat anti-rabbit Alexa 488 (1:400, #A-21422, 1:400, #A28175, ThermoFisher Scientific) in blocking solution for 2 h in RT. After the incubation, sections were rinsed 4 times for 10 min in PBS, then mounted and coverslipped with Vectashield + DAPI mounting medium (#H-1200, Vector Laboratories).

### Experiment 2 – Verification of virus injection-sites

The day after the last behavioral test we anesthetized the mice with a lethal dose of pentobarbital (Mebunat Vet 60 mg/ml) and transcardially perfused them with ice cold PBS followed by ice cold 4% PFA in PBS, followed by 24 h post-fixation in +4°C. Sagittal sections were cut using a cryostat (Leica RM2235 microtome, Leica Biosystems) at 35 μm. Serial sections were washed with PBS and mounted with ProLong Diamond hardset mounting medium. The innate fluorescence of the eGFP and mCherry were detected using 3DHISTECH Pannoramic 250 FLASH II digital slide scanner at Genome Biology Unit supported by HiLIFE and the Faculty of Medicine, University of Helsinki, and Biocenter Finland. Mice with bilateral expression of eGFP in the mPFC and mCherry expression in the ventral hippocampus (including CA1/3 subregions) were included in the analysis.

### Experiment 2 – Nodes of Ranvier immunohistochemistry

Free-floating sagittal sections were rinsed 3 times for 10 minutes in PBS, followed by incubation for 1h at RT in blocking solution (5% NGS, 2.5% BSA, 0.25% Triton X-100 in PBS). After blocking, the sections were incubated overnight in +4°C with anti-mouse CASPR (1:500, #75-001, Neuromab) in blocking solution. This was followed by rinsing 4 times for 10 min in PBS, and incubation with goat anti-mouse Alexa 647 (1:400, #ab150115, Abcam) in blocking solution for 2h at RT. Thereafter, sections were mounted and coverslipped with mounting medium (Immuno-mount, Thermoscientific).

### Imaging

#### Experiment 1

Imaging was performed with ZEISS LSM 880 Confocal Laser Scanning microscope with AiryScan (Zeiss). The distance from the bregma and the position of the ACC (layer 5/6) or forceps minor for each section was first determined at 10X magnification with a mouse brain atlas^56^. Nodes were then identified with a 63X oil objective in layers V/VI of the ACC and in the forceps minor. To image individual nodes within a field of view, a region around a node was cropped, and a z-stack of the cropped region was acquired. Z-stacks were acquired at a resolution of 0.04 x 0.04 x 0.10 μm.

#### Experiment 2

Imaging was performed as above but the hippocampal fimbria was first identified using 20X magnification. Paranodes, identified with 63X oil objective, overlapping with mCherry axons (mCherry+) as well as paranodes that did not co-localize with mCherry (mCherry-) were imaged within the hippocampal fimbria.

### 3D segmentation and morphometry of paranodes and juxtaparanodes

We developed an automated pipeline to segment and analyze the morphology of paranodes and juxtaparanodes, as well as to measure the length of nodes of Ranvier in the acquired 3D microscopy images. The pipeline initially segmented paranodes and juxtaparanodes applying geometric deformable models. However, because more than one pair of paranodes or juxtaparanodes were captured in the acquired images, the pipeline determined the main orientation of the segmented paranodes and juxtaparanodes and excluded those not along the main orientation, as shown in Supplementary Figure 4 a-f.

In more detail, we first applied a 3D median filter using a 5 x 5 x 3 sliding window to denoise the acquired 3D images of paranodes (red channel) and juxtaparanodes (green channel) separately to each channel. For segmentation we fused the median-filtered red and green channels into a single channel 3D image, denoted as *I*, by taking the maximum intensity value between the two channels at each voxel (Supplementary **Figure 4a**). To segment the 3D image *I*, first, we applied Frangi filtering^57^ to *I* to enhance its curvilinear structures, i.e., paranodes and juxtaparanodes, and suppress the background. Then, we thresholded the enhanced image to generate a 3D binary image used to initialize the Chan-Vese active surface model^58^. We used the implementation of the Chan-Vese model available in Matlab’s Image Processing Toolbox (version 2018b). We set the parameters as follows: contraction bias was 0.1, smoothness factor was 0.1, and the maximum number of iterations was 100. Applying the connected component analysis to the segmentation result, we generated a preliminary segmentation of paranodes and juxtaparanodes denoted as *L* (Supplementary **Figure 4b**). To exclude segmented components other than the paranodes and juxtaparanodes of interest, we first generated a 2D maximum intensity projection of the label image *L* along the direction of the focal plane, z-axis, as shown in Supplementary **Figure 4c**. We applied Hough transform^59^ to the maximum projection image of *L* to detect line segments in the image. We used the slope of the longest detected line segment, which represented the main orientation of the segmented paranodes and juxtaparanodes, to draw a line *l*\* that expanded to the image borders. The dashed line in Supplementary Figure 4c shows *l\** in the maximum projection image of *L* associated with the main orientation of the segmented components. Then each segmented component was projected on *l*\*, and its projection length was measured (Supplementary **Figure 4d**). The segmented components associated with the two longest projections, with non-intersecting projections, were selected as the final labels for the paranodes and juxtaparanodes of interest. For that, we first selected the longest projection and then the second longest projection that did not intersect with the longest projection. Because we applied the segmentation on the fused image, we used the two final segmented components as the initialization surfaces to segment paranodes and juxtaparanodes on 3D median-filtered images separately, using the Chan-Vese model with the same parameter settings as described earlier. Supplementary Figures 4e and f show the segmentation boundary of the paranode and juxtaparanode of interest in their corresponding channels.

We quantified morphological aspects of the segmented paranodes and juxtaparanodes in 3D following the approach in references ^60,61^. We first extracted the skeleton of paranodes by applying a distance transform-based skeletonization method from^62^ (Supplementary **Figure 4g_1_** and **g_2_**). With a plane perpendicular to the skeleton, we automatically extracted cross-sections along the length of segmented paranodes. The cross-sectional morphology of paranodes was quantified by the equivalent diameter and the length of the minor and major axes of the fitted ellipse. Moreover, we measured the length of paranodes by measuring the arc length of the acquired skeletons, as shown in Supplementary **Figure 4g**. The same procedures were applied to analyze the morphology of juxtaparanodes. Denote the set of voxel coordinates in two distinct paranodes by *A* and *B.* We measured the length of a node of Ranvier, Supplementary Figure 4g_2_, by using a robust version of 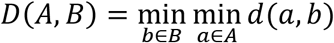, where *d*(.) is the Euclidean distance between two points.

Define the distance between the set of points *S* and a point *r* as 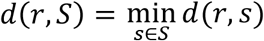. Then, the robust distance between two paranodes was defined as follows:

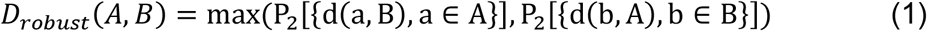

where *P*_2_ is the 2^nd^ percentile. In addition, we reported the intersection volume between the paranodes and juxtaparanodes.

### Statistical analysis

We assessed group differences in node and paranode morphology with generalized estimating equations (GEEs) to control for within-subject dependencies of individual nodes sampled from the same animal. This approach was selected to capture the within subject variance. GEE is suited for analyzing data with non-independent features^50,63^. For analyzing individual paranodes belonging to the same node, we also included node as a within-subjects factor (paranodes flanking the same node are presumed to be non-independent). Pairwise contrasts were computed to compare stress groups within strains (Con vs Res, Con vs Sus and Res vs Sus) with Fisher’s LSD, with manual test-wise Bonferroni correction (Experiment 1 α = 0.0167; Experiment 2 α = 0.0125). We compared behavioral test differences between groups using an unpaired (two-tailed) Student’s t test or a Mann-Whitney U test in case data were non-normally distributed.

## Acknowledgments

This work was funded by the Academy of Finland (316282; to IH; 323385; to AS), European Research Council Starting Grant (GenAnx; to IH), Sigrid Jusélius Foundation (to IH), and Finnish Cultural Foundation (to M-KK). We thank Kai Kaila and Hovatta lab members for helpful discussions and comments on the manuscript. Imaging was performed at the Biomedicum Imaging Unit and Genome Biology Unit and behavioral analyses at the Mouse Behavioural Phenotyping Facility, all supported by the Helsinki Institute of Life Science (HiLIFE) and Biocenter Finland.

## Author contributions

M-K.K., M.A.L., I.H. designed the experiments, interpreted results, and wrote the manuscript. M-K.K. performed immunohistochemical studies, microscopy imaging and data analysis with the assistance of S.H.J. and V.A. M.A.L. performed the CSDS experiment, DREADD manipulation, behavioral testing, immunohistochemical and data analysis with the help of G.M. and P.H. A.A. performed 3D reconstructions under the supervision of A.S. and J.T. who established the method. A.G. and K.T. performed gene expression and GSEA data analyses. All of the authors commented on the manuscript.

## Data availability

Gene expression data is available from the Gene Expression Omnibus with accession number GSE109315. The other data is available from the corresponding author upon reasonable request.

---

**Supplementary figure 1.**
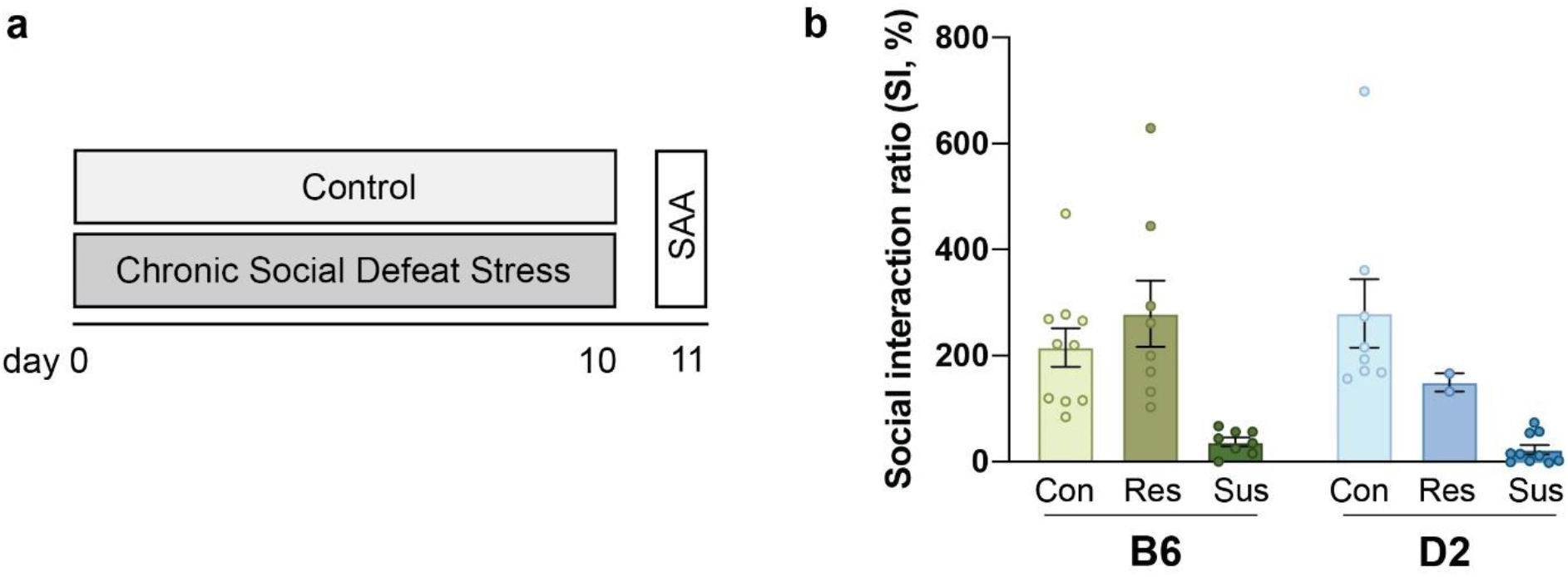
Grouping of mice to stress susceptible and resilient based on social interaction ratio. **a** Experimental timeline. **b** Grouping of mice into stress-resilient and -susceptible individuals, similar as in^50^. SAA=Social approach-avoidance test. Error bars represent ±SEM.

**Supplementary figure 2.**
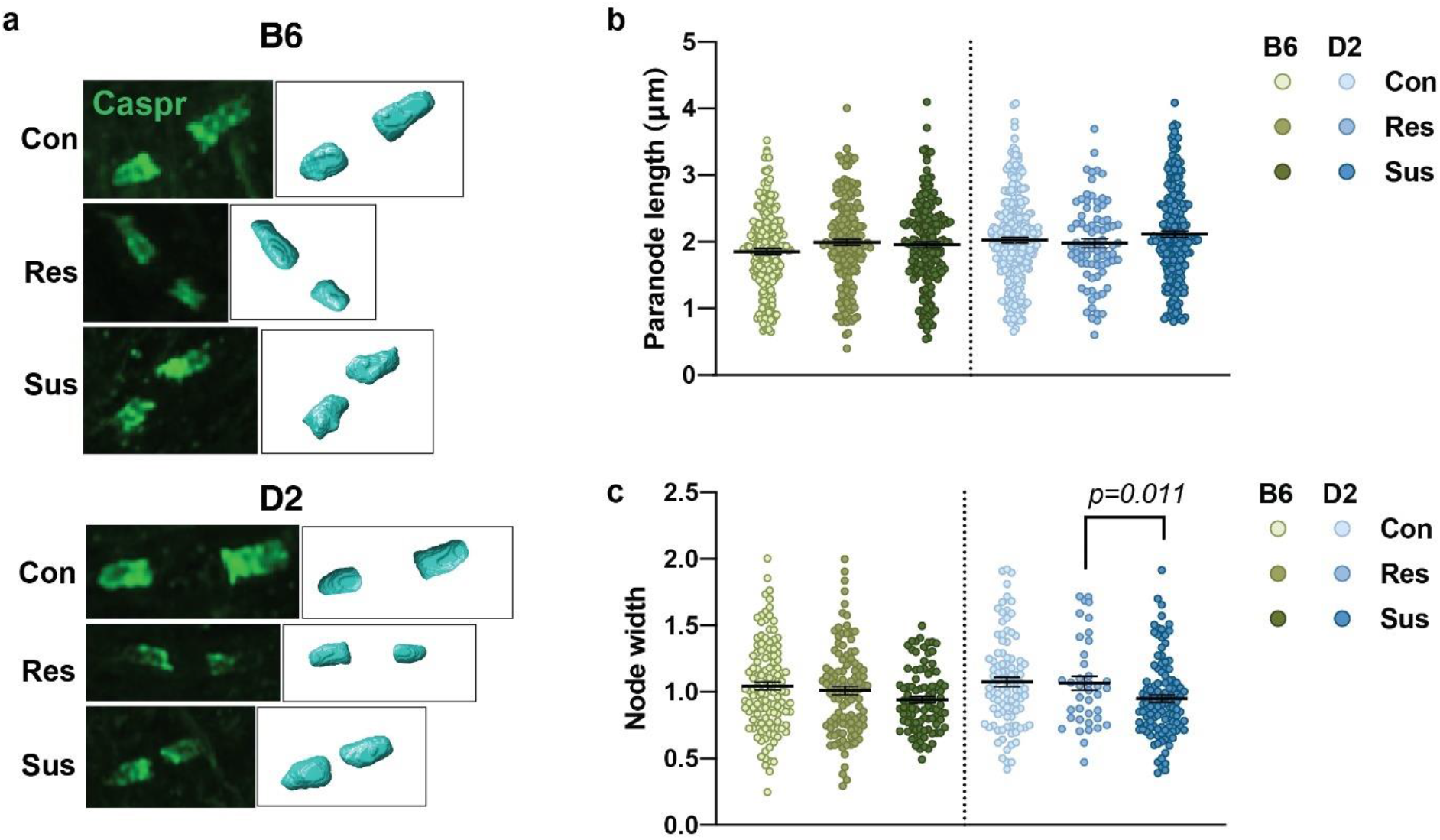
Shorter white matter node gaps in stress-susceptible D2 mice. **a** 3D reconstruction of paranodes in the forceps minor. **b,c** Quantification of paranode length and (**c**) node width in the white matter. (**b**B6: Con=234 nodes from 8 mice, Res=197 nodes from 6 mice, Sus=188 nodes from 5 mice; Generalized estimating equation (GEE), *p*=0.056. D2: Con=272 nodes from 6 mice, Res=80 nodes from 3 mice, Sus=219 nodes from 6 mice; GEE, *p*=0.252. **c** B6: Con=122 nodes from 8 mice, Res=110 nodes from 9 mice, Sus=94 nodes from 5 mice; GEE, *p*=0.164. D2: Con=136 nodes from 6 mice, Res=39 nodes from 3 mice, Sus=112 nodes from 6 mice; GEE, *p*=0.037, pairwise comparisons: Con *vs*. Res, *p*=0.465; Con *vs*. Sus, *p*=0.336; Res *vs*. Sus, *p*=0.011). Error bars represent ±SEM. Nominal *p*-values surviving Bonferroni correction are shown.

**Supplementary figure 3.**
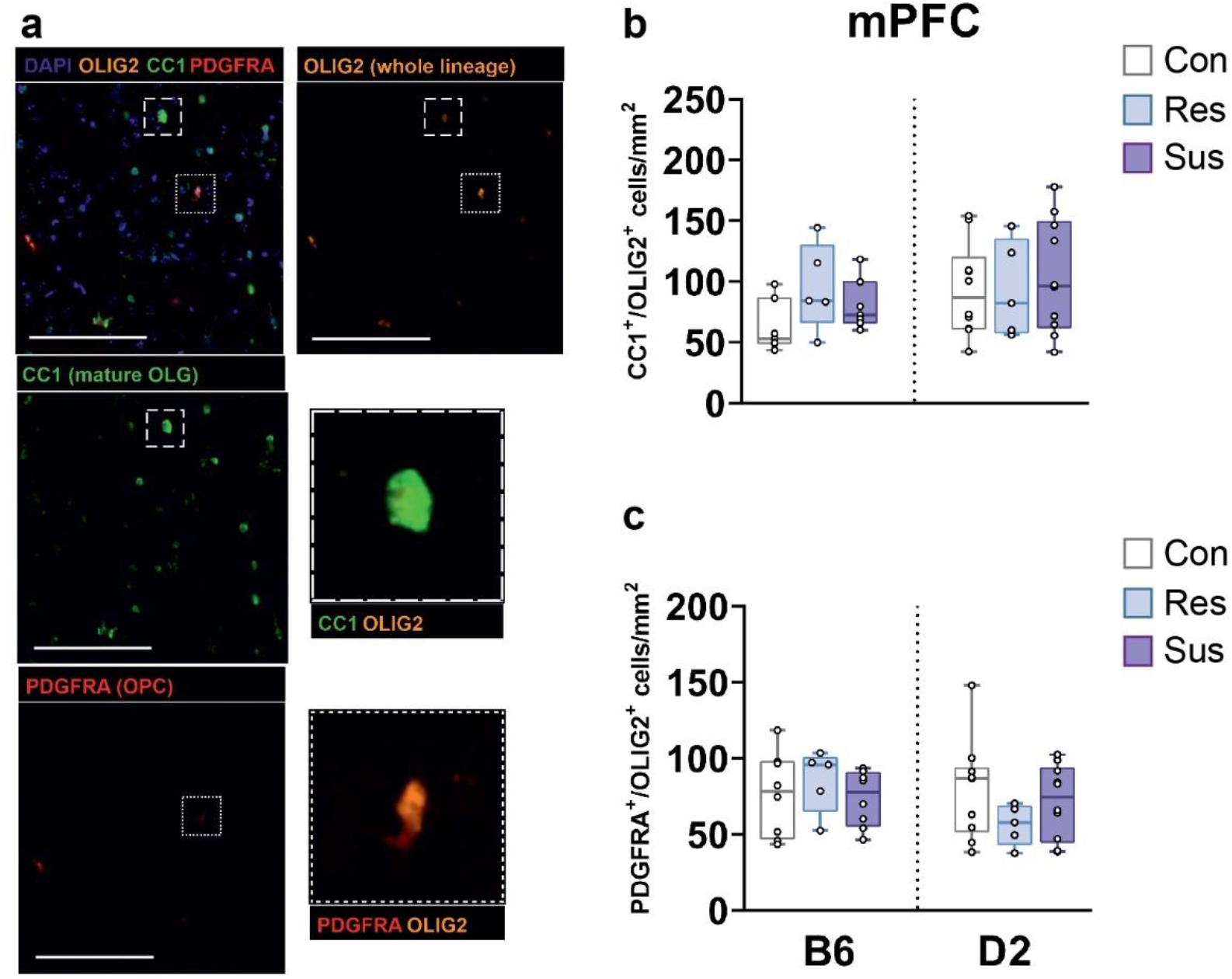
No difference in the density of oligodendrocyte lineage cells 1 week after chronic social defeat stress. **a** Oligodendrocyte precursor cells (OPCs) identified with platelet-derived growth factor receptor α (PDGFRA) and differentiated oligodendrocytes (OLGs) labeled with adenomateous polyposis coli clone 1 (CC1) antibody. These markers were co-stained with OLIG2 to restrict the analysis to OLG lineage. **b-c** Box plots of OLGs (**b**) and OPCs (**c**) in the mPFC. Each data point represents one animal’s average number of cells across 3-14 images.

**Supplementary figure 4.**
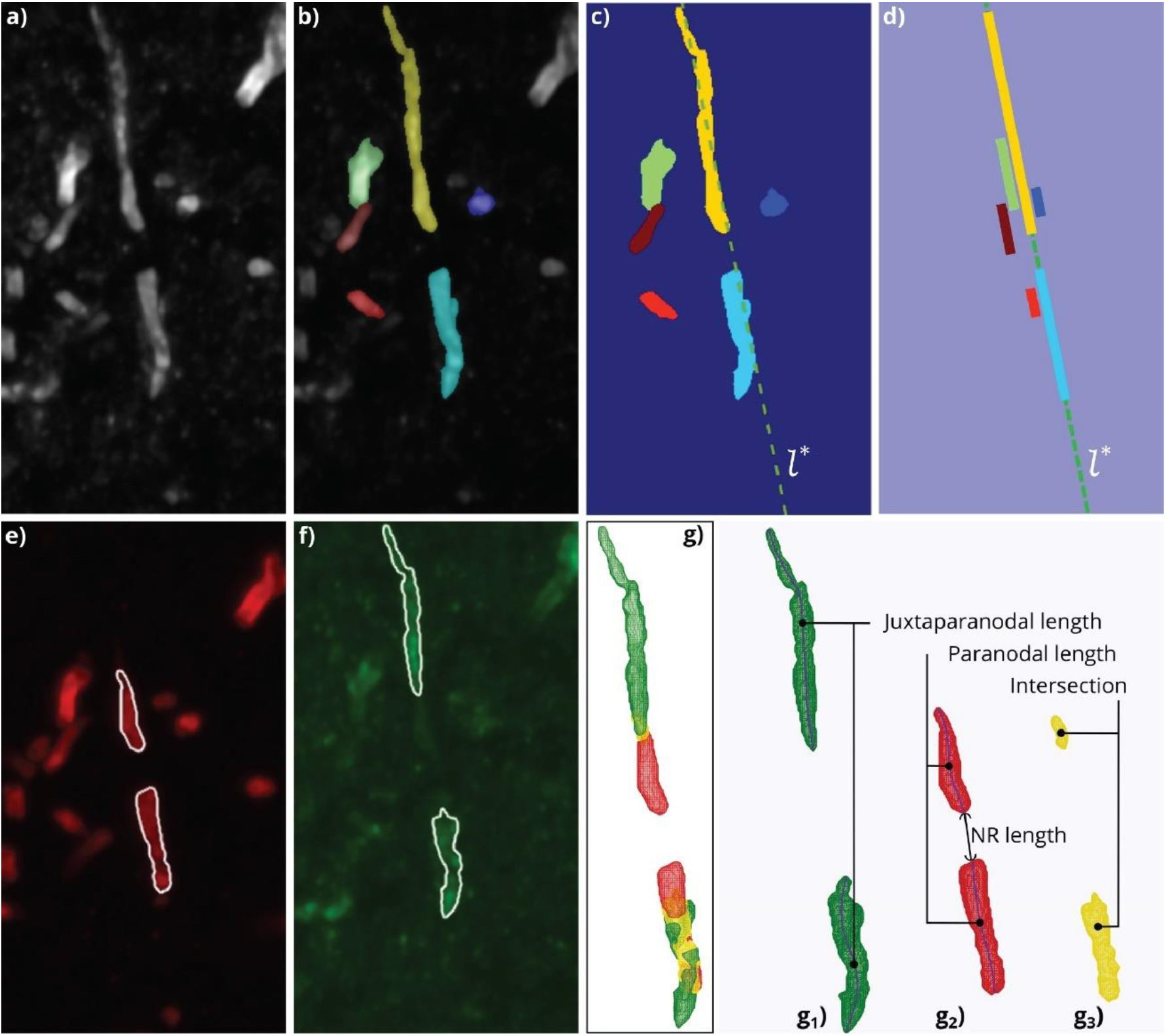
Automated segmentation and 3D morphometry of paranodes and juxtaparanodes in confocal microscopy images. **a**) A maximum projection image of paranodes (red channel) and juxtaparanodes (green channel). **b**) A preliminary segmentation of paranodes and juxtaparanodes using the Chan-Vese active surface model. **c**) The slope of the longest detected line segment, which associates with the main orientation of the segmented components, was used to draw a line *l*\* (dashed line) that expanded to the image borders. **d**) Projection of the segmented components on *l*\*. Colors correspond with the segmented components in panel c. **e**) Segmentation of paranodes visualized on the maximum projection image of the red channel. **f**) Segmentation of juxtaparanodes visualized on the maximum projection image of the green channel. **g**) 3D rendering of juxtaparanodes (green, g_1_, paranodes (red, g_2_, and their intersection (yellow, g_3_). The extracted skeletons of the paranodes and juxtaparanodes are overlaid on their corresponding rendering. NR=node of Ranvier.

**Supplementary table 1.**
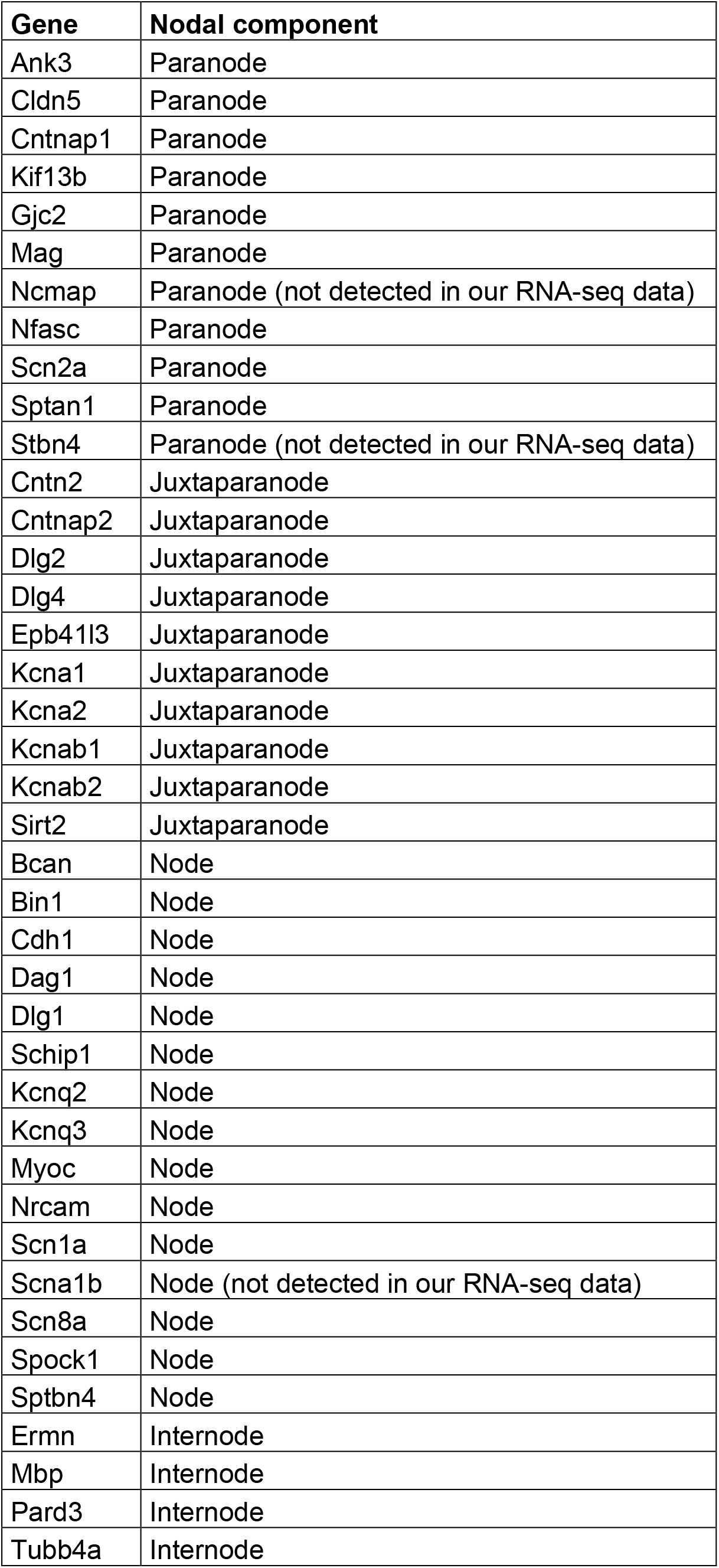
List of Node of Ranvier genes

